# Cell re-entry assays do not support models of pathogen-independent translocation of AvrM and AVR3a effectors into plant cells

**DOI:** 10.1101/038232

**Authors:** Benjamin Petre, Michaela Kopischke, Alexandre Evrard, Silke Robatzek, Sophien Kamoun

## Abstract

The cell re-entry assay is widely used to evaluate pathogen effector protein uptake into plant cells. The assay is based on the premise that effector proteins secreted out of a leaf cell would translocate back into the cytosol of the same cell via a yet unknown host-derived uptake mechanism. Here, we critically assess this assay by expressing domains of the effector proteins AvrM-A of *Melampsora lini* and AVR3a of *Phytophthora infestans* fused to a signal peptide and fluorescent proteins in *Nicotiana benthamiana*. We found that the secreted fusion proteins do not re-enter plant cells from the apoplast and that the assay is prone to false-positives. We therefore emit a cautionary note on the use of the cell re-entry assay for protein trafficking studies.

---

Pathogens deliver virulence proteins known as effectors into host tissues to promote compatibility (Win *et al*., 2012). How effectors enter host cells is a fundamental question in plant pathology that remains poorly understood (Ellis *et al*., 2006; Petre and Kamoun, 2014a). In filamentous plant pathogens (fungi and oomycetes), a current model postulates that effectors carry N-terminal entry domains (or translocation signals) that are necessary and sufficient to enter plant cells (Whisson *et al*., 2007; Dou *et al*., 2008; Rafiqi *et al*., 2010; Kale *et al*., 2010; Ve *et al*., 2013). Two assays are commonly used to evaluate the ability of effector proteins to enter host cells and are known as the ‘protein uptake’ and the ‘cell re-entry’ assays (Catanzariti *et al*., 2006; Dou *et al*., 2008; Kale and Tyler, 2011). In the protein uptake assay, purified effectors are directly applied to plant cells and evaluated for translocation. In the cell re-entry assay, effector proteins carrying an N-terminal secretory signal peptide are expressed in plant cells and secreted into the apoplast from where they re-enter the cell. In both assays, effectors encode an avirulence activity (AVR) or are fused to a fluorescent protein tag. In the case of AVR effectors, the readout for cell entry is hypersensitive cell death, a well-known response triggered upon activation of a corresponding cytosolic immune receptor. With fluorescent protein tags, the readout for cell entry is the visualization of the fluorescent signal inside the cell by confocal microscopy.

The protein uptake and the cell re-entry assays are increasingly used in the field of effector biology, yet their validity is frequently debated (http://tinyurl.com/qgqkv7b; Petre and Kamoun, 2014a). For instance, the robustness of the protein uptake assay has been critically evaluated and discussed (Wawra *et al*., 2013; Tyler *et al*., 2013). Whereas Tyler and colleagues concluded that the assay is robust and specific, Wawra and colleagues challenged this conclusion based on the observation that fluorescent proteins enter plant cells in a non-specific manner. The cell re-entry assay, which has been used over the last 10 years in several prominent studies (Supplementary Table 1), has also been criticized for lack of conclusiveness (Bos *et al*., 2006; Oh *et al*., 2009). Indeed, in this assay it is unclear whether proteins that accumulate in the cytosol have re-entered the plant cell from the apoplast, escaped from the secretory pathway through retrograde transport, or underwent translation at alternative start sites that resulted in proteins lacking a signal peptide. To date a critical evaluation of the cell re-entry assay has not been reported.

The effector proteins AvrM-A and AVR3a were used in landmark effector trafficking studies (Whisson *et al*., 2007; Rafiqi *et al*., 2010). In both proteins, ‘translocation’ or ‘cell entry’ domains have been defined, in which particular amino acid residues are required for cell entry (Whisson *et al*., 2007; Ve *et al*., 2013). AvrM-A is a 343-amino-acid secreted protein from the flax rust fungus *Melampsora lini*, which carries a 50-amino-acid entry domain within its N-terminal region (Rafiqi *et al*., 2010; Ve *et al*., 2013; Figure 1A). AVR3a is a 147-amino-acid secreted protein from the oomycete *Phytophthora infestans*, also known as the Irish potato famine pathogen (Armstrong *et al*., 2005; Bos *et al*., 2006). AVR3a carries a 37-amino-acid RXLR domain downstream of its signal peptide that is required for translocation inside host cells (Whisson *et al*., 2007; Figure 1A). In addition, the RXLR domain of AVR3A is similar in sequence to the RXLR domain of the effector Avr1b from *Phytophthora sojae*, which was described to mediate entry into host cells in the absence of the pathogen (Dou *et al*., 2008; Kale *et al*., 2010). Given the number of studies on translocation of AvrM-A, AVR3A, and related proteins into plant cells, we selected these two effectors to assess the robustness of the cell re-entry assay.

**Figure 1.**
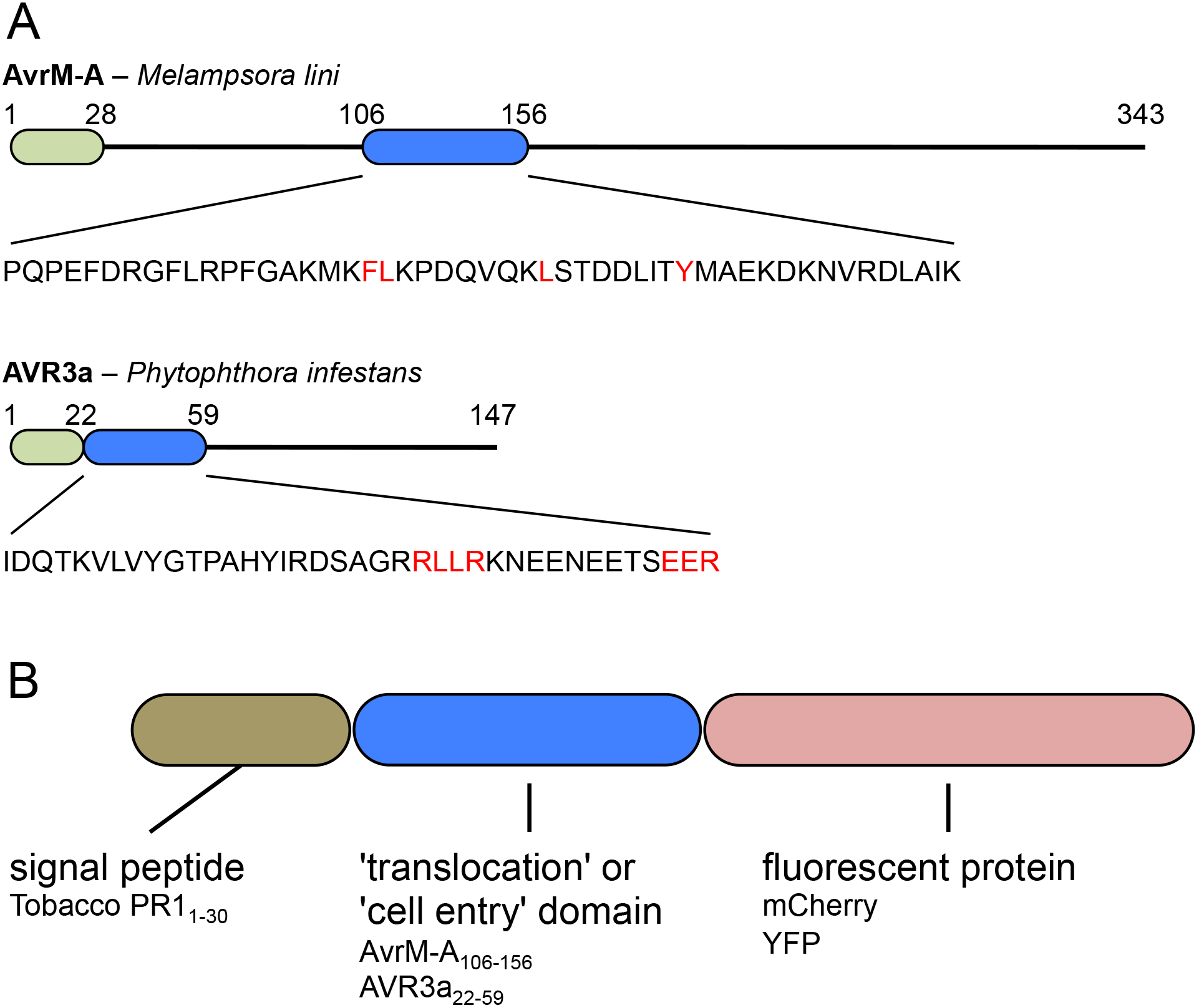
Sequences and constructs used in this study. **(A)** Schematic representation of AvrM-A and AVR3a effector proteins. Signal peptide for secretion (light green); reported translocation signal or uptake domain (blue); numbers: amino acid positions. Amino acids in red have been reported to be critical for cell entry activity, in these proteins or in similar proteins. **(B)** Schematic representation of the fusion proteins used in this study.

We first generated plant cell expression vectors to express chimeric proteins consisting of the previously defined translocation domains of AvrM-A or AVR3a fused to mCherry (a monomeric variant of the red fluorescent protein) targeted to the apoplast by the signal peptide of the tobacco Pathogenesis-related Protein 1 (PR1) (Figure 1B). Given that the PR1 signal peptide is highly effective in targeting proteins to the apoplast via the Endoplasmic Reticulum (ER)/Golgi secretory pathway (Hammond-Kosack *et al*., 1994), we expected the fusion proteins to accumulate in the apoplast. We then hypothesized that extracellular fusion proteins will be taken up by leaf cells as previously proposed (Supplementary Table 1). To challenge this hypothesis, we aimed at imaging leaf cells that do not express the fusion proteins but are surrounded by cells secreting the fusion proteins. This is possible because of two features of *Nicotiana benthamiana* transient transformation mediated by *Agrobacterium tumefaciens* (so-called “agroinfiltration” assay). First, infiltration of *A. tumefaciens* can produce mosaic transformation resulting in non-transformed leaf pavement cells that are surrounded by transformed cells. Second, *A. tumefaciens* infiltration of *N. benthamiana* leaves does not transform the guard cells, which are typically surrounded by transformed leaf pavement cells (Petre and Kamoun, 2014b). To this end, we expressed the PR1_1-30_-AvrM-A_106-156_-mCherry and PR1_1-30_-AVR3a_22-59_-mCherry fusions in *N. benthamiana* leaves and imaged non-transformed pavement and guard cells whose immediate apoplast was saturated with fusion proteins. As illustrated in Figure 2, we observed multiple cases of non-transformed pavement cells (n>30) without any fluorescent signal, indicating that they did not take up the extracellular fusion proteins. In addition, all the guard cells we observed (n>200) lacked any fluorescent signal indicating that the apoplastic fusion proteins did not enter neighboring guard cells (Figure 2, Supplementary Figure 1). Plasmolysis experiments confirmed the accumulation of the fusion proteins in the apoplast (Figure 3). We confirmed that the non-transformed cells we observed appear healthy and integral, based on the absence of autofluorescence, occurrence of mobile and evenly distributed chloroplasts, and a steady cytosolic stream. We conclude that under these conditions the entry domains of AvrM-A and AVR3a do not mediate translocation of the fusion proteins from the apoplast into *N. benthamiana* leaf cells.

**Figure 2.**
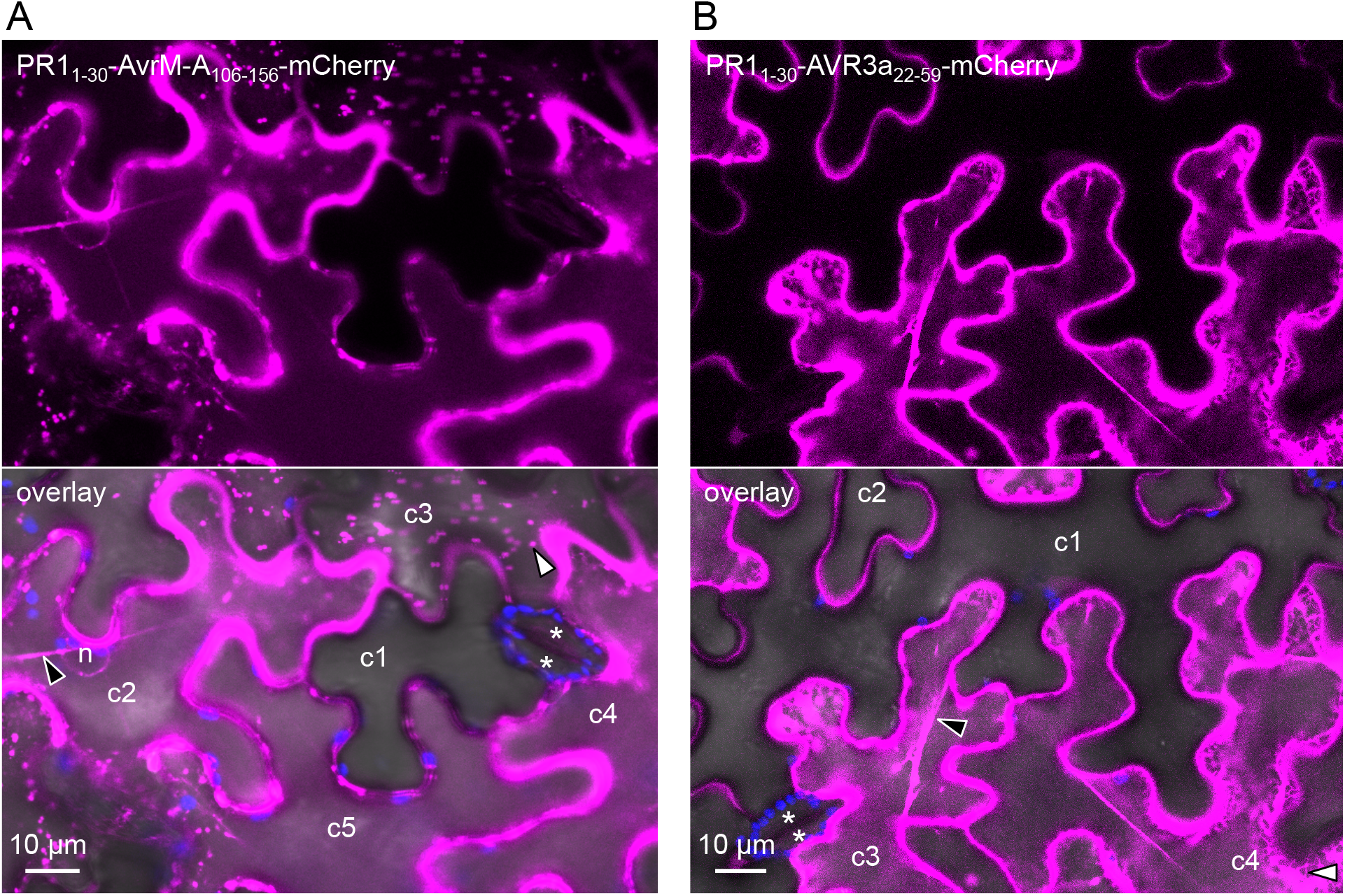
Leaf cells do not take up extracellular AvrM-A- and AVR3a-mCherry fusions. **(A)** Live-cell imaging of PR1_1-30_-AvrM-A_106-156_-mCherry in *Nicotiana benthamiana* leaves. Images show a single optical section. Cells marked c1 to c5 are pavement cells differentially accumulating the fusion proteins (c1: no accumulation; c2 to c5: high accumulation). White arrowhead: mCherry-labelled puncta; black arrowhead: mCherry-labelled cytosol. n: nucleus. Asterisks: guard cells. Note the absence of mCherry signal in c1 and in guard cells. **(B)** Live-cell imaging of PRl_1.30_-AVR3a_22.59_-mCherry in *N. benthamiana* leaves. Images show a maximal projection of three optical sections (z-stack: 2.4 μm). Cells marked c1 to c4 are pavement cells differentially accumulating the fusion proteins (c1 and c2: no accumulation; c3 and c4: high accumulation). White arrowhead: mCherry-labelled stuctures resembling the endoplasmic reticulum; black arrowhead: mCherry-labelled cytosol. n: nucleus. Note the absence of mCherry signal in c1, in c2, and in guard cells.

In addition to the apoplast, we observed accumulation of the AvrM-A and AVR3a mCherry fusion proteins inside transformed cells (Figure 2 and 3). We aimed at better determining the intracellular accumulation pattern of these proteins, but the fluorescent signal from the apoplast interfered with – and impaired optimal imaging of – the intracellular signal. To circumvent this issue, we constructed new fusion proteins using the Yellow Fluorescent Protein (YFP), a protein that does not fluoresce in acidic environments such as the apoplast (Shaner *et al*., 2005). Following infiltration of the produced *A. tumefaciens* strains in *N. benthamiana* leaves and live cell imaging, we noted that both PR1_1-30_-AvrM-A_106-156_-YFP and PR1_1-30_-AVR3a_22-59_-YFP fusions accumulated in Golgi bodies and the ER, respectively (Figure 4, Supplementary Figure 2). Interestingly, high accumulation in these secretory compartments correlated with the appearance of a weak fluorescent signal in the cytosol (Figures 2-4). These observations indicate that some fusion proteins end up in the cytosol of the cells in which they are expressed, and that this phenomenon correlates with high accumulation in the secretory pathway. Altogether, these data suggest that cytosolic accumulation could be a consequence of saturation of secretory compartments rather than cell re-entry, and is therefore a possible source of false positives.

**Figure 3.**
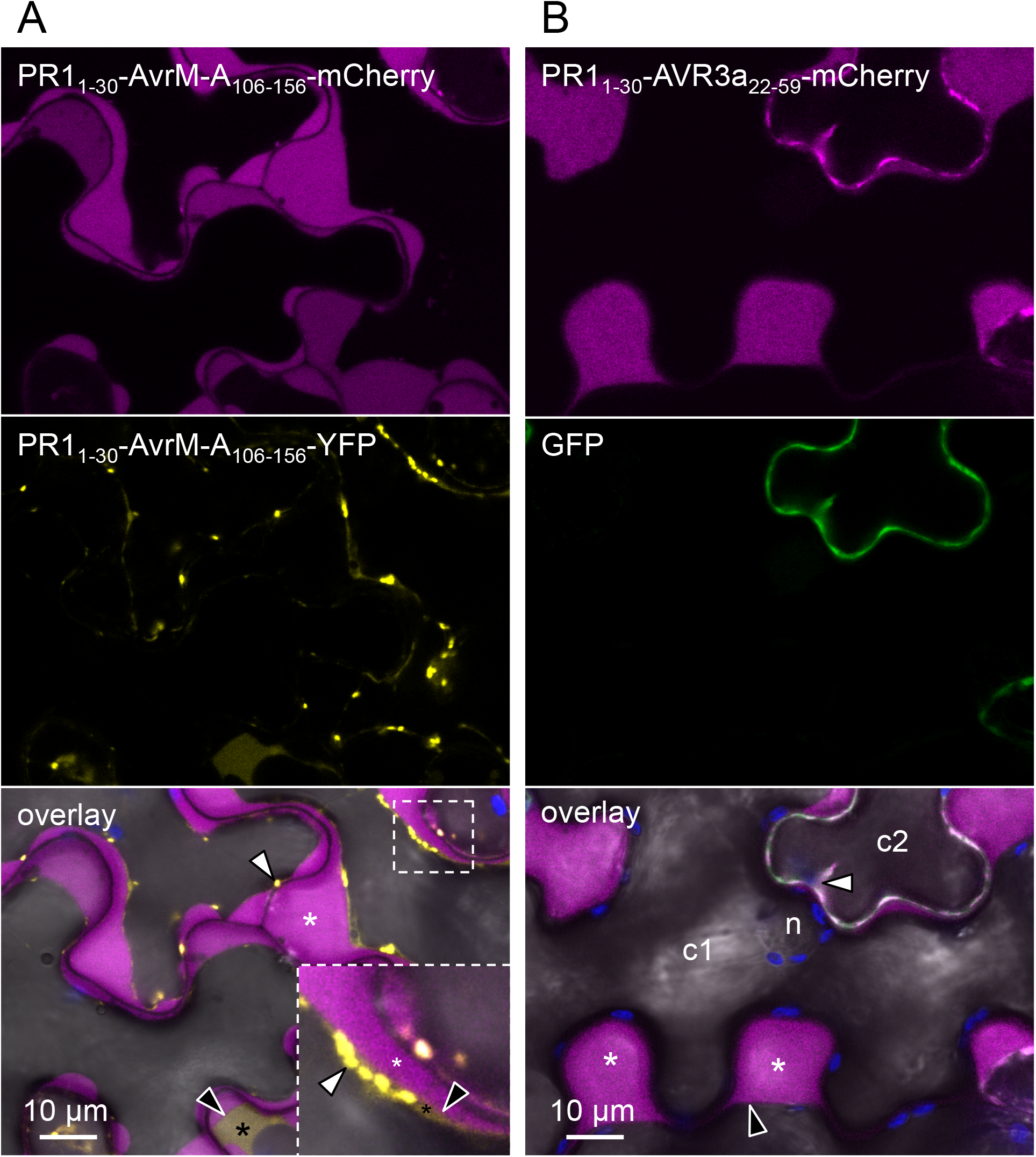
Secreted AvrM-A- and AVR3a-mCherry fusions accumulate in the apoplast. **(A)** Live-cell imaging of PRl_1–30_-AvrM-A_106–156_-mCherry and PR1_1–30_-AvrM-A_106–156_-Yellow Fluorescent Protein (YFP) in *Nicotiana benthamiana* leaves treated with 1 M NaCL for 5 minutes. Images show a single optical section. Black asterisk: cytosol; white arrowheads: YFP-labelled puncta. The insert in the bottom-right part of the overlay image shows a close-up of the white rectangle. **(B)** Live-cell imaging of PR1_1–3_-AVR3a_22–59_-mCherry and a green fluorescent protein (GFP, nucleus/cytosol marker) in *N. benthamiana* leaves. Images show a single optical section. Cells marked c1 and c2 are pavement cells differentially accumulating the proteins (c1: no accumulation; c2: high accumulation of both proteins). White arrowhead: cytosol; n: nucleus. White asterisks: apoplast; black arrowhead: plasma membrane separated from the cell wall.

**Figure 4.**
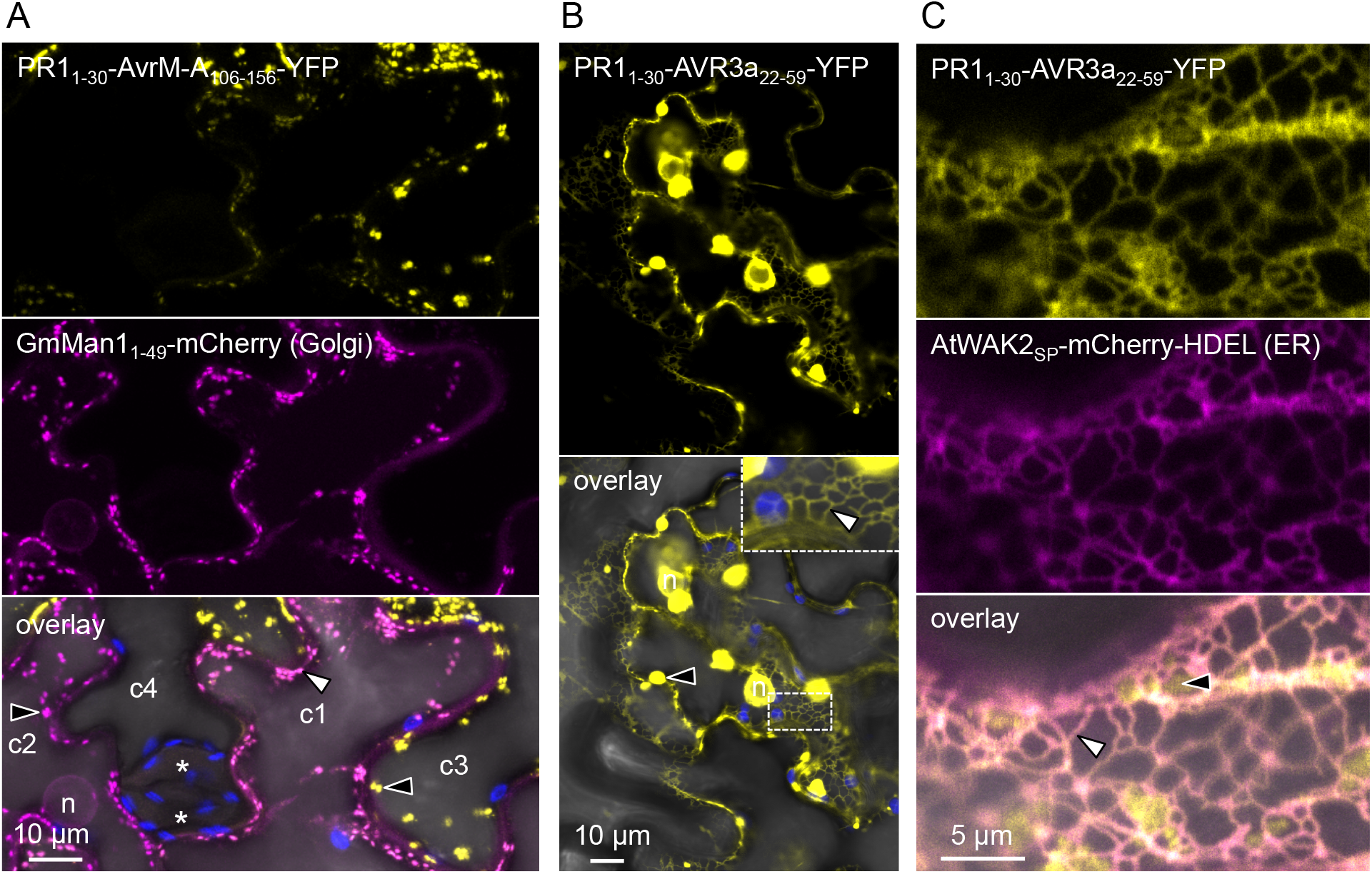
Secreted AvrM-A- and AVR3a-YFP fusions accumulate in the cytosol of the cells in which they are expressed. **(A)** Live-cell imaging of PR1_1-30_-AvrM-A_106-156_-Yellow Fluorescent Protein (YFP) and GmMan1_1-49_-mCherry (Golgi marker) in *Nicotiana benthamiana* leaf cells. Images show a maximal projection of seven optical sections (z-stack: 5.6 μm). Cells marked c1 to c4 are pavement cells differentially accumulating the fusion proteins (c1: high accumulation of both fusions; c2: high accumulation of GmManl_1-49_-mCherry; c3: high accumulation of PR1_1-30_-AvrM-A_106-156_-YFP; c4: no accumulation). Asterisks: guard cells; white arrowheads: Golgi bodies with overlapping YFP and mCherry signals; black arrowheads: Golgi bodies with mCherry (magenta in c2) or YFP (yellow in c3) signal. **(B)** Live-cell imaging of PR1_1-30_-AVR3a_22-59_-YFP in *N. benthamiana* leaf cells. Images show a single optical section. Black arrowheads: aggregates labelled by the YFP signal; white arrowheads: endoplasmic reticulum (ER)-like structures labelled by the YFP signal. The insert in the top-right part of the overlay image shows a close-up on the ER-like structures in the white rectangle. **(C)** Live-cell imaging of PR1_1-30_-AVR3a_22-59_-YFP and AtWAK2_SP_-mCherry-HDEL (ER marker) in *N. benthamiana* leaf cells. Images show a single optical section. Black arrowhead: YFP signal outside of the ER; white arrowhead: ER. YFP signal was collected at 530-580 nm. n: nucleus.

In summary, we failed to conclusively demonstrate cell uptake of the translocation domains of AvrM-A and AVR3A using live cell imaging of *N. benthamiana* leaves transformed with effector domains fused to fluorescent proteins. Therefore, the conclusion that these proteins can enter plant cells in the absence of the pathogen may need to be revisited. It also appears that a sub-population of proteins targeted to the secretory pathway accumulates in the cytosol and thus could be a source of false positives in the cell re-entry assay. We conclude that a positive result in the cell re-entry assay may not necessarily indicate uptake from the apoplast. We recommend a reinterpretation of the previous data generated with this assay and the associated models for effector translocation into plant cells.

## METHODS

### Biological material and cloning procedures

We built fusion proteins containing the ‘translocation sequence’ or ‘entry domain’ of AvrM-A (amino acid positions 106 to 156; AvrM-A_106-156_) or AVR3a (amino acid positions 22 to 59; AVR3A_22-59_), which would be secreted by plant cells and traceable by fluorescent microscopy (Figure 1A). To drive the secretion of the fusion proteins, we used the signal peptide of *Nicotiana tabacum* Pathogenesis-Related protein 1 (PR1) (amino acid positions 1 to 30; PR1_1-30_). We amplified the coding sequence the signal peptide of PR1 by polymerase chain reaction from a previously published plasmid (Kamoun *et al*., 1999). As fluorescent tags, we used the YFP and the mCherry. We obtained the coding sequences of AVR3a and AvrM-A through gene synthesis (Genewiz, London, UK). We used the Golden Gate technology to assemble DNA fragments and to build the four following chimera: PR1_1-30_-AvrM-A_106-156_-mCherry, PR1_1-30_-AvrM-A_106-156_-YFP, PR1_1-30_-AVR3A_22-59_-mCherry, and PR1_1-30_-AVR3A_22-59_-YFP (Figure 1B, Supplementary Table S2). The DNA fragments were individually assembled in the binary vector pICH86988, immediately downstream of a cauliflower mosaic virus 35S promoter, as previoulsy described (Petre *et al*., 2015). We used *Escherichia coli* (DH5α), *A. tumefaciens* GV3101 (pMP90), and *N. benthamiana* as previously described (Petre *et al*., 2015).

### Transformation of N. benthamiana leaf cells and laser-scanning confocal microscopy

We delivered T-DNA constructs into leaf cells of three-week-old *N. benthamiana* plants by infiltration of *A. tumefaciens*, following the method previously described (Win *et al*., 2011, Petre *et al*., 2015). To favor the mosaic transformation of leaf pavement cells by infiltration of *A. tumefaciens*, we used a bacterial OD_600_ of 0.1 and we collected leaves as soon as 36 hours post infiltration for immediate use. We observed the leaf tissues at the edge of the infiltrated leaf area, where heterogeneous transformation of leaf cells is more common.

We performed live-cell imaging with a Leica DM6000B/TCS SP5 laser-scanning confocal microscope (Leica microsystems, Bucks, UK), using a 63x (water immersion) objective as previously described (Petre *et al*., 2015). GFP, YFP and mCherry were excited using 488 nm, 514 nm, and 561 nm laser lines, respectively. We collected fluorescence emission between 505-525 nm for GFP, 680-700 nm for chlorophyll autofluorescence, 530-550 nm for YFP, and 580-620 nm for mCherry, except otherwise stated. We induced plasmolysis by incubating leaf tissues 5 minutes in 1 M NaCL. We performed image analysis with Fiji (http://fiji.sc/Fiji). We used the following markers for co-localisation experiments: GmMan1_1-49_-mCherry (Golgi), AtWAK2_SP_-mCherry-HDEL (ER), mRFP-ARA7 (early and late endosomes), and ARA6-mRFP (late endosomes and multi vesicular bodies) (Nelson *et al*., 2007). The FM4-64 dye (Synapto RED, 5 mg/mL in water, working solution 1:2000) was used to trace endocytic compartments (Bolte *et al*., 2004), following procedures previously described (Beck *et al*., 2012).

## ACKNOWLEDGMENTS

We thank K. Schumacher (Heidelberg, Germany) for providing plasmids. We thank M. Mbengue for discussions. BP was supported by an INRA Contrat Jeune Scientifique (CJS), by the European Union, in the framework of the Marie-Curie FP7 COFUND People Programme, through the award of an AgreenSkills’ fellowship (under grant agreement n° 267196). Research at The Sainsbury Laboratory is supported by the Gatsby Charitable Foundation and the BBSRC. MK is supported by a grant of the European Research Council (ERC) awarded to SR.

## AUTHOR CONTRIBUTIONS

BP and SK conceived the study and designed the experiments. BP, MK, and AE carried out the research. SR contributed to the design of experiments and provided expertise in cell biology. BP prepared the first draft of the manuscript. BP and SK contributed to the preparation of the manuscript. BP, MK, SR, and SK were involved in the revision of the manuscript. All authors have agreed to the final content.

---

**Supplementary Figure 1.**
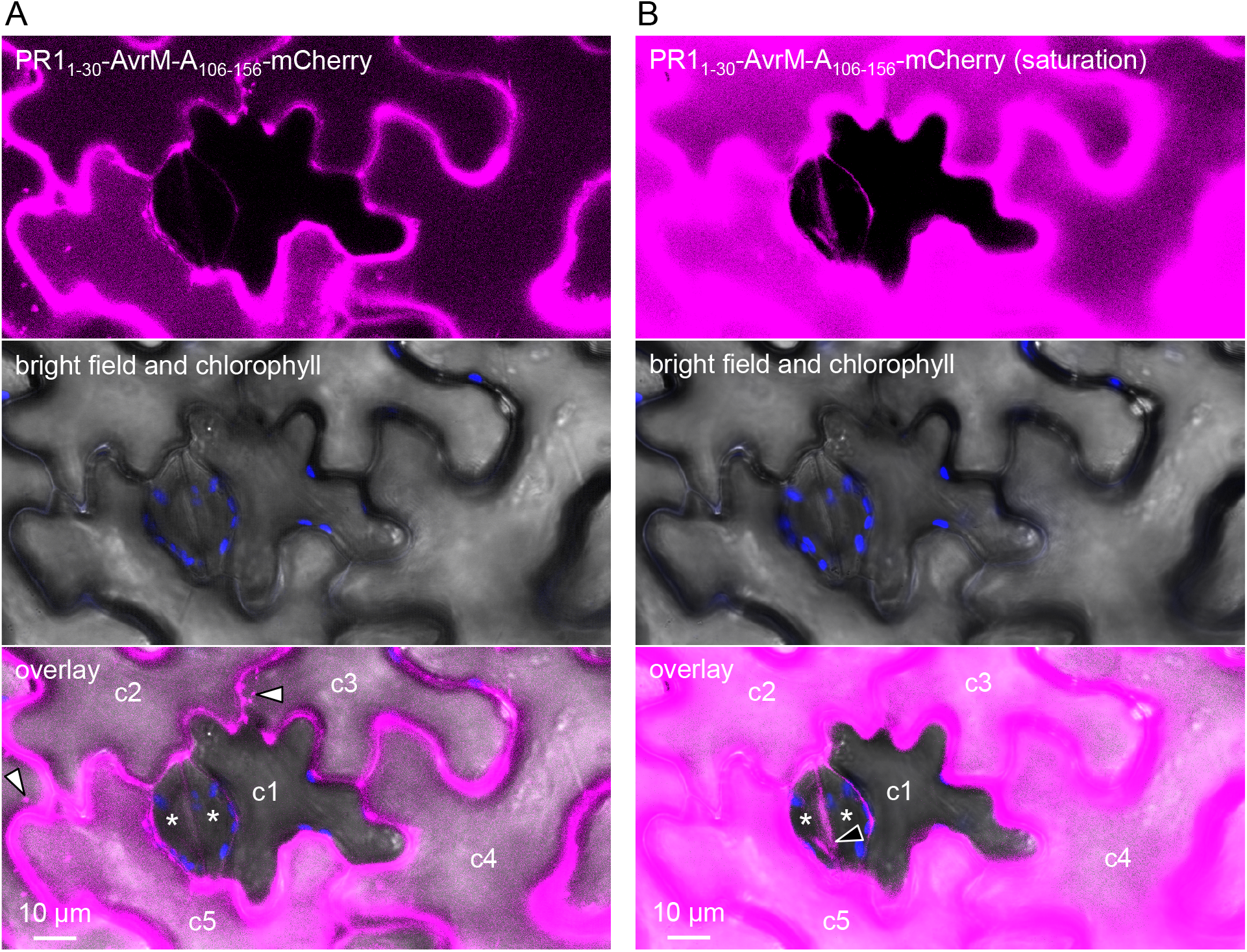
Non-transformed leaf cells do not exhibit fluorescent signal. **(A)** Live-cell imaging of PR1_1-30_-AvrM-A_106-156_-mCherry in *Nicotiana benthamiana* leaf cells. Images show a single optical section. Cells marked c1 to c5 are pavement cells differentially accumulating the fusion proteins (c1: no accumulation; c2 to c5: high accumulation). Asterisks: guard cells; white arrowheads: mCherry-labelled Golgi bodies. **(B)** Same as A with a higher laser power. Black arrowhead: autofluorescence of the ostiole rim. Note the absence of fluorescent signal in cl and in guard cells.

**Supplementary Figure 2.**
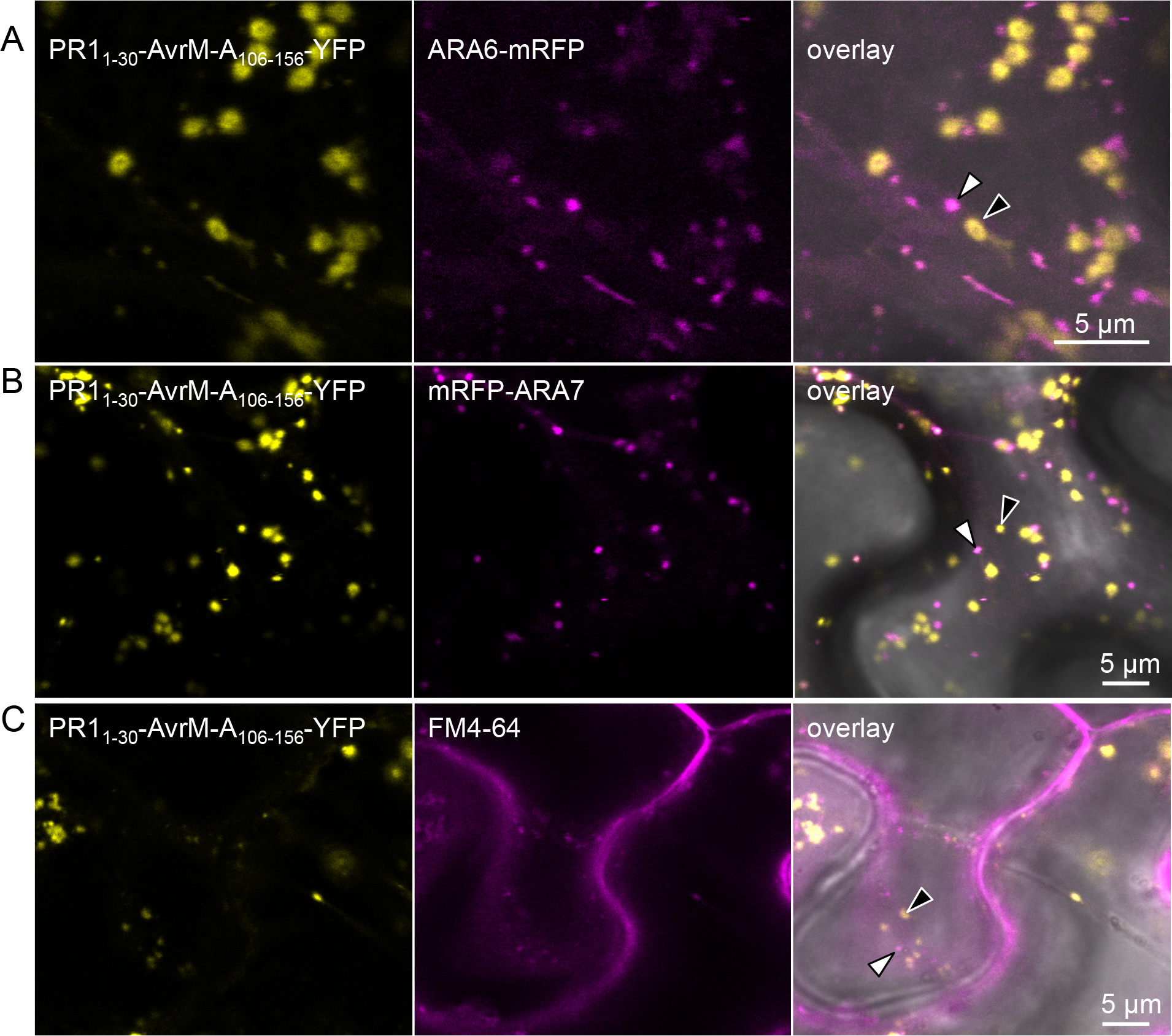
PR1_1-30_-AvrM-A_106-156_-YFP does not accumulate in ARA6-, ARA7-, and FM4-64-labelled vesicles. **(A)** Live-cell imaging of PR1_1-30_-AvrM-A_106-156_-Yellow Fluorescent Protein (YFP) and ARA6-monomeric red fluorescent protein (mRFP) (Late endosome [LE] and multi vesicular body [MVB] marker) in *Nicotiana benthamiana* leaf cells. Images show a single optical section. White arrowhead: LE/MVB; black arrowhead: YFP-labelled Golgi body. (B) Live-cell imaging of PR1_1-30_-AvrM-A_106-156_-YFP and mRFP-ARA7 (early endosome [EE] and LE marker) in *N. benthamiana* leaf cells. Images show a single optical section. White arrowhead: EE/LE; black arrowhead: YFP-labelled Golgi body.

**Supplementary Table 1.** List of studies that used cell re-entry assays

| Reference | plant species |
| --- | --- |
| Catanzariti *et al.*, 2006, Plant Cell | *N. benthamiana* |
| Bos *et al.*, 2006, Plant J | *N. benthamiana* |
| Dou *et al.*, 2008, Plant Cell | Soybean, Onion |
| Oh *et al.*, 2009, Plant Cell | *N. benthamiana* |
| Rafiqi *et al.*, 2010, Plant Cell | Flax, Tobacco |
| Kale *et al.*, 2010, Cell | Soybean |
| Gu *et al.*, 2011, PLOS ONE | Soybean |
| Ribot *et al.*, 2013, Plant J | Rice |
| Na *et al.*, 2013, Mol Plant Microbe Interact | Soybean |
| Ve *et al.*, 2013, Proc Natl Acad Sci U S A | Tobacco |

**Supplementary Table 2.**
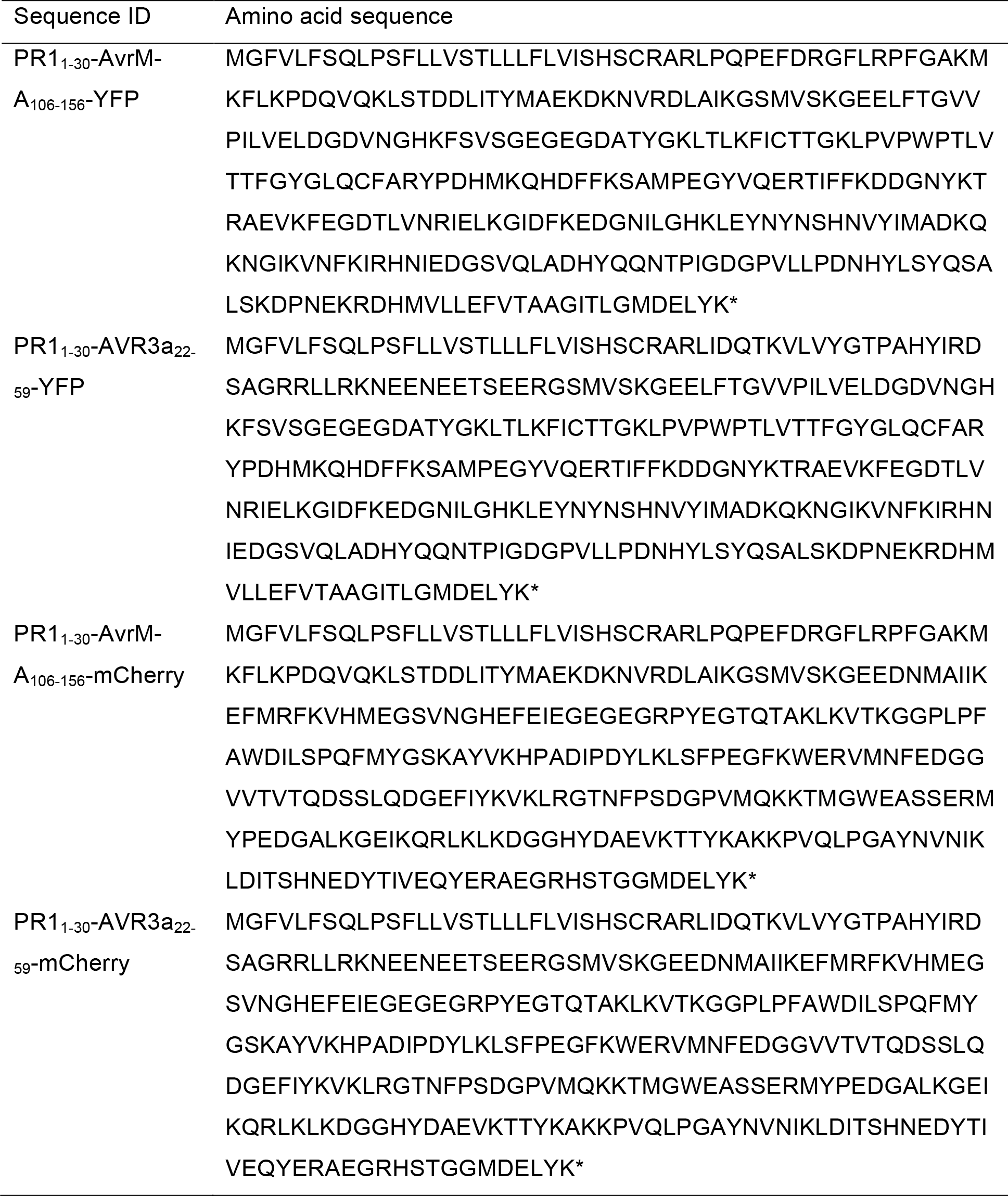
Sequences of the fusion proteins used in this study

## REFERENCES

Armstrong MR, Whisson SC, Pritchard L, Bos JIB, Venter E, Avrova AO, Rehmany AP, Böhme U, Brooks K, Cherevach I, Hamlin N, White B, Fraser A, Lord A, Quail MA, Churcher C, Hall N, Berriman M, Huang S, Kamoun S, Beynon JL, Birch PRJ. 2005. An ancestral oomycete locus contains late blight avirulence gene *Avr3a*, encoding a protein that is recognized in the host cytoplasm. Proc Natl Acad Sci U S A 102:7766–71

Beck M, Zhou J, Faulkner C, MacLean D, Robatzek S. 2012. Spatio-Temporal Cellular Dynamics of the Arabidopsis Flagellin Receptor Reveal Activation Status-Dependent Endosomal Sorting. Plant Cell 24:4205–19

Bolte S, Talbot C, Boutte Y, Catrice O, Read ND, Satiat-Jeunemaitre B. 2004. FM-dyes as experimental probes for dissecting vesicle trafficking in living plant cells. J Microsc 24:159–73

Bos JI, Kanneganti TD, Young C, Cakir C, Huitema E, Win J, Armstrong MR, Birch PR, Kamoun S. 2006. The C-terminal half of *Phytophthora infestans* RXLR effector AVR3a is sufficient to trigger R3a-mediated hypersensitivity and suppress INF1-induced cell death in *Nicotiana benthamiana*. Plant J 24:165–76

Catanzariti AM, Dodds PN, Lawrence GJ, Ayliffe MA, Ellis JG. 2006. Haustorially expressed secreted proteins from flax rust are highly enriched for avirulence elicitors. Plant Cell 24:243–56

Dou D, Kale SD, Wang X, Jiang RH, Bruce NA, Arredondo FD, Zhang X, Tyler BM. 2008. RXLR-mediated entry of *Phytophthora sojae* effector Avr1b into soybean cells does not require pathogen-encoded machinery. Plant Cell 24:1930–47

Ellis J, Catanzariti AM, Dodds P. 2006. The problem of how fungal and oomycete avirulence proteins enter plant cells. Trends Plant Sci 24:61–3

Gu B, Kale SD, Wang Q, Wang D, Pan Q, Cao H, Meng Y, Kang Z, Tyler BM, Shan W. 2011. Rust Secreted Protein Ps87 Is Conserved in Diverse Fungal Pathogens and Contains a RXLR-like Motif Sufficient for Translocation into Plant Cells. PLOS ONE 24:e27217

Hammond-Kosack KE, Harrison K, Jones JD. 1994. Developmentally regulated cell death on expression of the fungal avirulence gene Avr9 in tomato seedlings carrying the disease-resistance gene Cf-9. Proc Natl Acad Sci U S A 24:10445–9

Kale SD, Tyler BM. 2011. Entry of oomycete and fungal effectors into plant and animal host cells. Cell Microbiol. 24:1839–48

Kale SD, Gu B, Capelluto DG, Dou D, Feldman E, Rumore A, Arredondo FD, Hanlon R, Fudal I, Rouxel T, Lawrence CB, Shan W, Tyler BM. 2010. External lipid PI3P mediates entry of eukaryotic pathogen effectors into plant and animal host cells. Cell 24:284–95

Kamoun S, Honée G, Weide R, Laugé R, Kooman-Gersmann M, de Groot K, Govers F, de Wit PJGM. 1999. The fungal gene *Avr9* and the oomycete gene *inf1* confer avirulence to Potato Virus X on tobacco. Mol Plant Microbe Interact 24:459–62

Na R, Yu D, Qutob D, Zhao J, Gijzen M. 2013. Deletion of the *Phytophthora sojae* avirulence gene Avr1d causes gain of virulence on Rps1d. Mol Plant Microbe Interact 24:969–76

Nelson BK, Cai X, Nebenführ A. 2007. A multicolored set of *in vivo* organelle markers for co-localization studies in *Arabidopsis* and other plants. Plant J 24:1126–36

Oh SK, Young C, Lee M, Oliva R, Bozkurt TO, Cano LM, Win J, Bos JI, Liu HY, van Damme M, Morgan W, Choi D, van der Vossen EA, Vleeshouwers VG, Kamoun S. 2009. *In planta* expression screens of *Phytophthora infestans* RXLR effectors reveal diverse phenotypes, including activation of the *Solanum bulbocastanum* disease resistance protein Rpi-blb2. Plant Cell 24:2928–47

Petre B, Lorrain C, Saunders DG, Win J, Sklenar J, Duplessis S, Kamoun S. 2015. Rust fungal effectors mimic host transit peptides to translocate into chloroplasts. Cell Microbiol doi: 10.1111/cmi.12530

Petre B, Kamoun S. 2014a. How do filamentous pathogens deliver effector proteins into plant cells? PLoS Biol 24:e1001801

Petre B, Kamoun S. 2014b. *Agrobacterium tumefaciens* does not transform guard cells in *Nicotiana benthamiana*. Figshare http://dx.doi.org/10.6084/m9.figshare.1002071

Rafiqi M, Gan PH, Ravensdale M, Lawrence GJ, Ellis JG, Jones DA, Hardham AR, Dodds PN. 2010. Internalization of flax rust avirulence proteins into flax and tobacco cells can occur in the absence of the pathogen. Plant Cell 24:2017–32

Ribot C, Césari S, Abidi I, Chalvon V, Bournaud C, Vallet J, Lebrun MH, Morel JB, Kroj T. 2013. The *Magnaporthe oryzae* effector AVR1-CO39 is translocated into rice cells independently of a fungal-derived machinery. Plant J 24:1–12

Shaner NC, Steinbach PA, Tsien RY. 2005. A guide to choosing fluorescent proteins. Nat Methods 24:905–9

Tyler BM, Kale SD, Wang Q, Tao K, Clark HR, Drews K, Antignani V, Rumore A, Hayes T, Plett JM, Fudal I, Gu B, Chen Q, Affeldt KJ, Berthier E, Fischer GJ, Dou D, Shan W, Keller NP, Martin F, Rouxel T, Lawrence CB. 2013. Microbe-independent entry of oomycete RxLR effectors and fungal RxLR-like effectors into plant and animal cells is specific and reproducible. Mol Plant Microbe Interact 24:611–6

Ve T, Williams SJ, Catanzariti AM, Rafiqi M, Rahman M, Ellis JG, Hardham AR, Jones DA, Anderson PA, Dodds PN, Kobe B. 2013. Structures of the flax-rust effector AvrM reveal insights into the molecular basis of plant-cell entry and effector-triggered immunity. Proc Natl Acad Sci U S A 24:17594–9

Wawra S, Djamei A, Albert I, Nürnberger T, Kahmann R, van West P. 2013. *In vitro* translocation experiments with RxLR-reporter fusion proteins of Avr1b from *Phytophthora sojae* and AVR3a from *Phytophthora infestans* fail to demonstrate specific autonomous uptake in plant and animal cells. Mol Plant Microbe Interact 24:528–36

Whisson SC, Boevink PC, Moleleki L, Avrova AO, Morales JG, Gilroy EM, Armstrong MR, Grouffaud S, van West P, Chapman S, Hein I, Toth IK, Pritchard L, Birch PR. 2007. A translocation signal for delivery of oomycete effector proteins into host plant cells. Nature 24:115–8

Win J, Chaparro-Garcia A, Belhaj K, Saunders DG, Yoshida K, Dong S, Schornack S, Zipfel C, Robatzek S, Hogenhout SA, Kamoun S. 2012. Effector biology of plant-associated organisms: concepts and perspectives. Cold Spring Harb Symp Quant Biol 24:235–47

Win J, Kamoun S, Jones AM. 2011. Purification of effector-target protein complexes via transient expression in *Nicotiana benthamiana*. Methods Mol Biol 24:181–94

